# Ultra long-lived plasma cells in the human small intestine produce microbiota-reactive IgA antibodies

**DOI:** 10.1101/2025.03.17.643697

**Authors:** Niladri Bhusan Pati, Brian K Chung, Kristian Holm, Frank Sætre, Henrik Mikael Reims, Anders Theodor Aasebo, Tobias Gedde-Dahl, Diana Domanska, Johannes R Hov, Espen Sønderaal Bækkevold, Frode Lars Jahnsen

## Abstract

A large fraction of the intestinal microbiota is highly coated with secretory IgA, and bacteria-specific IgA is believed to shape the composition of the microbiota. A hallmark of the adaptive immune system is immunological memory to specific antigens. However, whether there is strong and persistent memory of secretory antibodies to bacterial antigens has not been determined. Here we show that ultra long-lived CD19^-^ CD45^-^ (age>20 years) plasma cells (PCs) residing in the human small intestine produce IgA that binds to most taxa of a diverse anaerobic microbiota culture. Long-lived CD19^-^CD45^+^ (age>10 years) and short-lived CD19+CD45+ (age<2 years) PCs also produced IgA with broad bacterial reactivity. A clear correlation between high-binding and low-binding taxa was observed across the PC subsets. We also found that host PCs were depleted in acute intestinal graft versus host disease, a condition strongly associated with loss of microbiota diversity. Together, we show that bacterial antigens in the intestine induce an extremely stable, long-lasting humoral immune memory that may be important for the long-term stability and resilience of the intestinal microbiome.

## Introduction

Approximately 80% of all plasma cells (PCs) in the body are located in the intestinal tract producing several grams of antibodies every day^1^. Intestinal PCs produce mainly dimers and some larger polymers of IgA, whereas a minor fraction produces pentameric IgM. Both antibody classes interact with the transmembrane poly Ig receptor (pIgR) located on the basolateral side of secretory epithelial cells that actively translocate IgA and IgM to the apical side. The pIgR further proteolytically cleaved to release secretory Igs that perform their protective functions as free antibodies in the gut lumen^2^. Secretory IgA plays an important protective role in the defense against pathogens and toxins through a variety of non-inflammatory activities that increase their clearance and prevent their access to the intestinal epithelium. Moreover, IgA can reduce bacterial virulence and growth^3,4,5^.

A large fraction of the intestinal microbiota is coated with IgA (and IgM)^16^ and several lines of evidence suggest that antibodies generated in response to microbiota colonization shape the gut microbiota composition. In humans, IgA deficient individuals exhibit gut microbiota dysbiosis despite secretion of compensatory IgM^7,8^. Moreover, patients with common variable immunodeficiency (CVID), the most common symptomatic primary immunodeficiency that is characterized by the lack of IgG in addition to IgA and/or IgM, show extensive microbial alterations in the gut, including low microbial diversity, systemic inflammation, and gut leakage^9^. In addition, mice lacking B cells, IgA, Ig class switching or pIgR expression have a disturbed microbiota composition^10,11,12,13^.

Studies in mice have indicated that under homeostatic conditions the gut microbiota generates short-lived PCs with little somatic hypermutation that secrete polyreactive IgA antibodies with low microbial affinity, whereas mucosal pathogens and vaccines elicit high-affinity, T-cell dependent antibody responses^14^. However, studies also reveal that under homeostatic conditions, affinity maturation occur in Peyer’s patch germinal centers to obtain persistent gut antigen specific response^15,16^. Further, a recent study in humans showed that high microbiota reactivity of IgA requires somatic hypermutations^17^. Interestingly, several monoclonal antibodies cloned from human IgA+ intestinal PCs bound several phylogenetically diverse bacterial species (termed cross-species reactivity), that were unrelated to polyreactivity^17^. To reconcile these apparently contradictory findings it is possible that newborns may rely on “innate-like” polyreactive antibodies to shape a protective microbiota, whereas when the mucosal immune system develop with age, affinity-matured high somatic hypermutation antibodies with cross-species reactivity will dominate.

Studies in humans have shown that the gut microbiota composition is remarkably stable over time^18,19^. Moreover, antibiotic treatment and fecal transplantation, which dramatically change the microbiota composition, often appear to have only transient effects with gradual normalization towards baseline over time^20,21,22^. Interestingly to this end, we have demonstrated that the human small intestine (SI) contains a large fraction of PCs that survives for decades^23^. Given the importance of IgA in shaping the microbiota composition, it is tempting to speculate that these ultra long-lived PCs play an important role for long-term stability and resilience of the microbiota^24^. However, whether IgA secreted from ultra long-lived PCs are directed towards the microbiota has not been determined. To probe this, we harvested IgA produced by three subsets of SI PCs with very different lifespan^23^ and examined the IgA reactivity to a diverse culture of a human intestinal microbiome (ACHIM, Anaerobic Cultivated Human Intestinal Microbiota)^25^. Furthermore, we analyzed the replacement kinetics of host PCs in the gut mucosa of stem cell transplanted patients developing graft versus host disease.

## Results

### ACHIM constitutes a diverse culture of human intestinal microbiota

Experiments that examine IgA binding to the human gut microbiota often isolate bacteria from stool samples^6,17,26^. However, this approach is complicated by several factors. First, bacteria obtained from stool are coated with endogenous antibodies that need to be removed before use^17,26,27^. Secondly, culturing representative microbiomes that lack endogenous coating under anaerobic conditions has proven to be difficult^28^. Finally, the microbiota composition varies substantially between individuals and results using stool samples from different individuals are difficult to compare^18,29^.

To circumvent the challenges of using patient stool samples, here we used ACHIM, a human intestinal microbiota isolated from a healthy individual that has been successfully cultured under anaerobic conditions for many years^25^. ACHIM has been shown to have therapeutic effects on *Clostridium difficile* infections^30^ and is currently being tested in several clinical trials^25,31^. To determine the bacterial composition of ACHIM we performed 16S ribosomal RNA (rRNA) sequencing and showed that ACHIM contains the four main phyla of known commensal intestinal bacteria (*Firmicutes, Bacteroidota, Actinobacteriodota, and Proteobacteria*), covering 33 genera (Fig. 1A-B). Thus, ACHIM is a diverse culture of anaerobic microbiota that allowed us to analyze, under standardized conditions, microbiota binding of IgA produced by PCs isolated from the SI.

**Figure 1.**
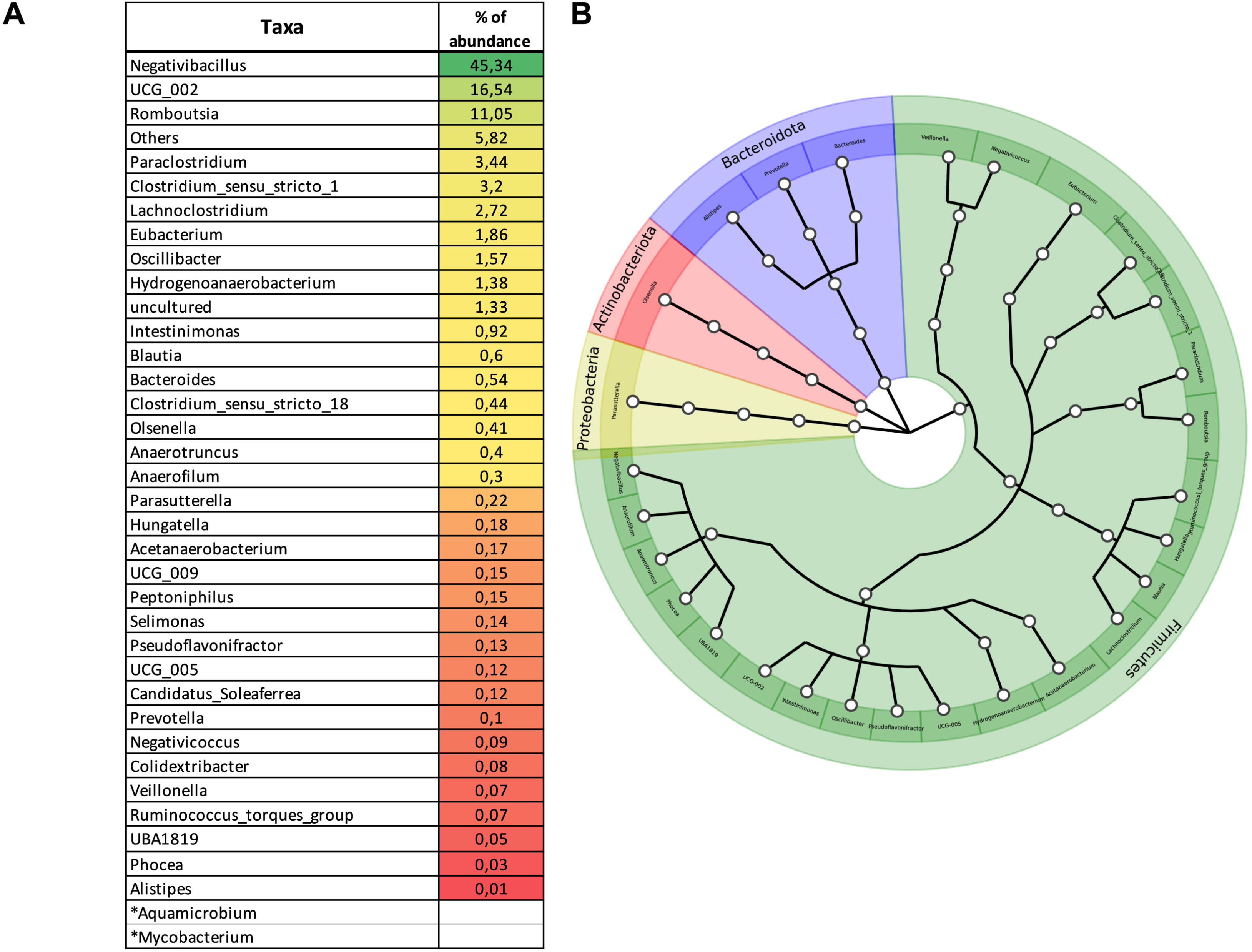
Representation of individual taxa in post sort ACHIM. **(A)** Shows representation of individual taxa and their percentage of relative abundance in ACHIM using 16S rRNA sequencing. * Indicates contaminants and are identified by comparing the 16s sequences between ACHIM that has passed through the FACS sorter and that does not. **(B)** Individual taxa are organized as a cladogram using Graphlan^57^ in an order of phylum to genus (annotations displayed in colors).

### Ultra long-lived and long-lived intestinal PCs produce microbiota-reactive IgA

To obtain PCs for ex vivo culture, we acquired distal duodenum-proximal jejunum resections during surgery of pancreatic cancer patients (n=4; Whipple procedure). The tissue was treated with collagenase to obtain single cell suspensions and PC subsets comprising short-lived CD19+CD45+ (double positive, DP) PCs, long-lived CD19-CD45+ (single positive, SP) PCs, and ultra long-lived CD19-CD45- (double negative, DN) PCs were isolated by magnetic bead-based separation as previously described^23^ (Fig. 2A). The purity of the isolated PC subsets was assessed with flow cytometry (Fig. 2B)^24^.

**Figure 2.**
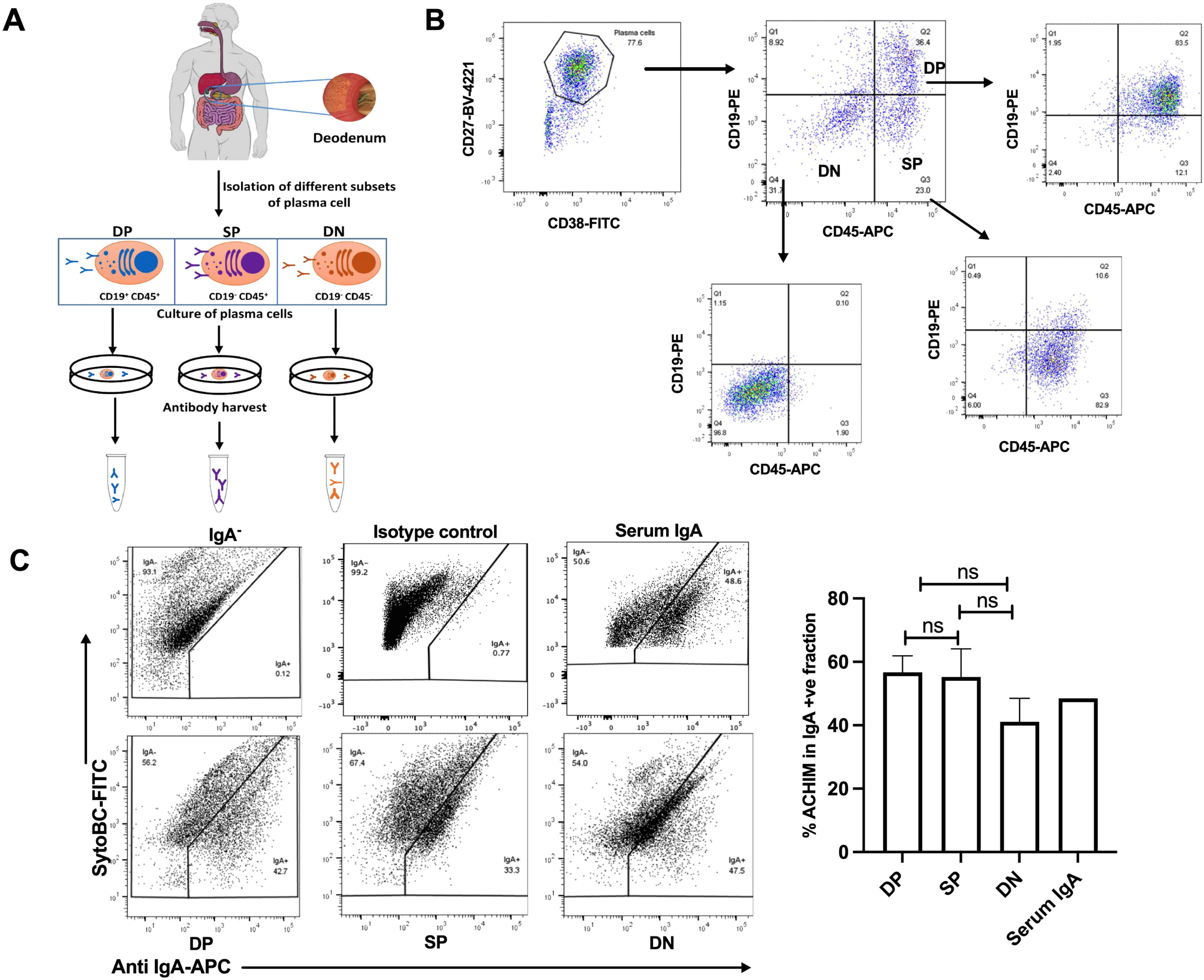
Culture of human intestinal plasma cells, antibody harvest and their binding affinity towards gut microbiota. **(A)** Schematic representation of isolation and culture of PCs from human small intestine for harvesting antibodies from the respective PC subsets. **(B)** FACS plots represents for the purity of PC subsets isolated from the small intestine resections (n=4) obtained during the Whipple procedure. **(C)** ACHIM, either unstained (negative control) or incubated with antibodies harvested from human intestinal PCs and purified IgA from human serum (positive control). The incubated bacteria are stained with secondary anti IgA-APC and sorted in the flow-cytometer as IgA^+^ and IgA^-^ fraction. Shown FACS plots are the representative of four individual experiments and the column graph shows the percentage of ACHIM that binds to IgA isolated from different PCs (n=4) or serum.

To determine if short-, long- and ultra long-lived PCs produced microbiota-reactive antibodies, sorted PC subsets were cultured for 48 hours and IgA-containing supernatants were collected. We incubated ACHIM (20x10^6^ bacteria) with the collected supernatant containing IgA (10 µg/ml), followed by staining with APC-conjugated anti-human IgA, and sorted bacteria into IgA^+^ and IgA^-^ fractions using FACS sorter. IgA from short-lived and long-lived PCs coated about 55% of the bacteria, whereas IgA from ultra long-lived PCs coated approximately 40% (Fig. 2C; Fig. S1). Serum IgA was included as positive control (Fig 2C).

To characterize the bacterial reactivity of IgA we performed 16S rRNA-seq of bacteria from the sorted IgA^+^ and IgA^-^ fractions. We first examined the richness within each sample (alpha diversity) and quantified differences in the overall taxonomic composition between samples (beta diversity). All samples showed high richness scores that were similar to pure ACHIM (Fig. 3A). Moreover, beta diversity showed no clear clustering of the samples; nor between the IgA+ or IgA-samples within each individual or between individuals (Fig. 3B).

**Figure 3.**
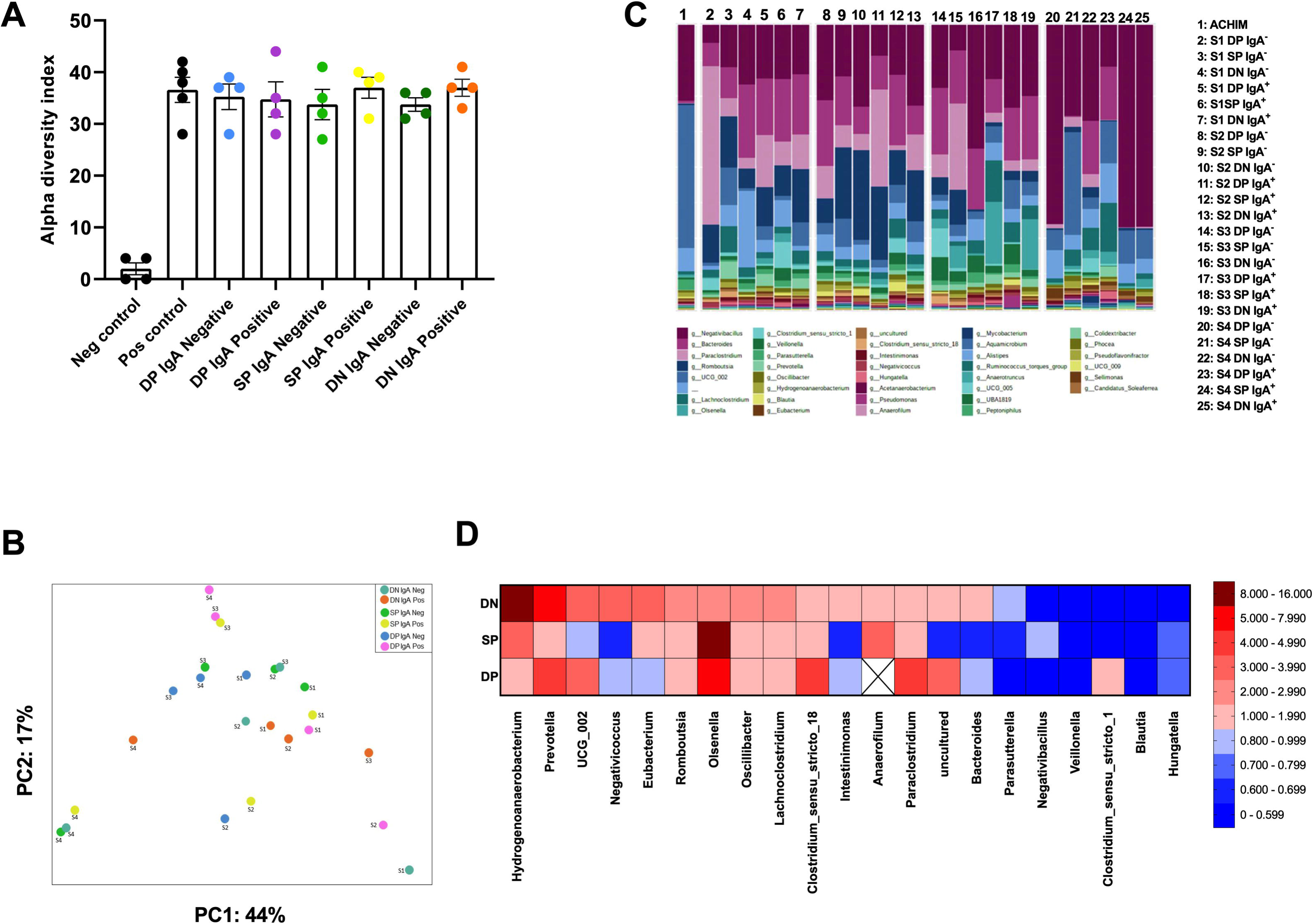
Human intestinal IgA predominantly targets and binds to diverse commensal bacteria. (A and B) α diversity and β-diversity parameters for sorted IgA^+^ and IgA^-^ ACHIM. The α-diversity indices are obtained by comparing the groups using one-way ANOVA and the β-diversity is measured using Bray-Curtis dissimilarity measure. (C) Average relative abundance of taxa in indicated 25 different fractions assessed by 16S sequencing. ‘ S’ represents to the patient sample. DN= double negative, SP= single positive and DP= double positive plasma cells. (D) Median value (n=4) plotted for the enrichment of individual taxa in the IgA+ fraction. DN, SP and DP represents to the plasma cells from where the IgA isolated.

To identify which bacterial taxa that was highly coated with IgA, we calculated the fraction of IgA+ bacteria taxa as a fraction of IgA-bacteria taxa with an enrichment score of >1 indicating a relative enrichment of the taxon in the IgA+ fraction.

Bacterial taxa with low representation in ACHIM (Fig. 1) were excluded as too few taxon counts were present in either the IgA+ or the IgA-fraction. IgA derived from all the PC subsets bound a broad range of taxa (Fig. 3C), and IgA derived from ultra long-lived PCs showed an enrichment score >1 in 15 out of 21 (70%) bacterial taxa (Fig. 3D). In comparison IgA from long-lived PCs showed an enrichment score >1 in 47% and short-lived PCs in 55% of bacterial taxa, respectively (Fig. 3D). Our findings thus indicated that intestinal IgA responses to the gut microbiota are extremely persistent.

We found a clear correlation between high binding and low binding bacteria across all PC subsets (Figure 3D). Focusing on the taxa that were highly coated by IgA from ultra long-lived PCs we found the highest enrichment score for Hydrogenanaerobacterium and Prevotella in all three PC subsets (Fig. 3D). Interestingly, Prevotella has previously been shown to be highly coated in various mouse models and shown to drive intestinal inflammation and disease^32,33,34,35,36^. Among bacteria with low IgA-binding reactivity, *Blautia and Veillonella* showed low IgA coating across all PC subsets (Fig. 3D). In contrast to Prevotella, these low binding bacteria, these low binding bacteria have been shown to have probiotic functions^37,38^.

### Intestinal PCs are rapidly depleted in acute gut graft versus host disease (GVHD)

As we observed that resident intestinal PCs possess antibody reactivities highly directed towards the gut microbiome, we sought to assess whether these properties play a role for the high resilience and stability observed in the human microbiota^18,39^. To investigate this possibility further we examined the association between microbiota diversity and the replacement kinetic rate of intestinal PCs in a human setting. Loss of microbiota diversity is strongly associated with the occurrence of acute intestinal GVHD in patients receiving allogeneic human stem cell transplantation (allo-HCT)^40,41,42^. We therefore hypothesized that there is a higher turnover of intestinal PCs in allo-HCT patients that develop acute gut GVHD compared to those that do not. To test this hypothesis, we analyzed the replacement kinetics of host PCs in allo-HCT patients with and without acute gut GVHD. We included male patients receiving female donor cells (n=16) that were biopsied on suspicion of acute GVHD. Clinical characteristics given in Table 1. Histological evaluation by an experienced pathologist showed that 11 of the patients had acute GVHD, whereas the other five patients did not show any histological signs of intestinal pathology. We performed immunofluorescence stainings in situ combining an antibody to the pan-PC marker CD38 with XY-FISH^43,44^, in which host PCs were identified as Y+CD38+ cells (Fig 4A; Fig. S2). All biopsies were obtained between day 20 and day 50 after transplantation and there was no significant change in the total number of PCs over time and no difference in the numbers of PCs between patients with and without acute GVHD (Fig. 4B, C). However, patients with acute GVHD had significantly lower precentage of host Y+CD38+ PCs than those without acute GVHD (Fig. 4D). This was independent of the preconditioning regime (Fig. 4E). Together, these findings showed that intestinal PCs are more rapidly replaced when acute GVHD developed, which strengthen our notion that PC longevity is related to microbiota diversity.

**Figure 4.**
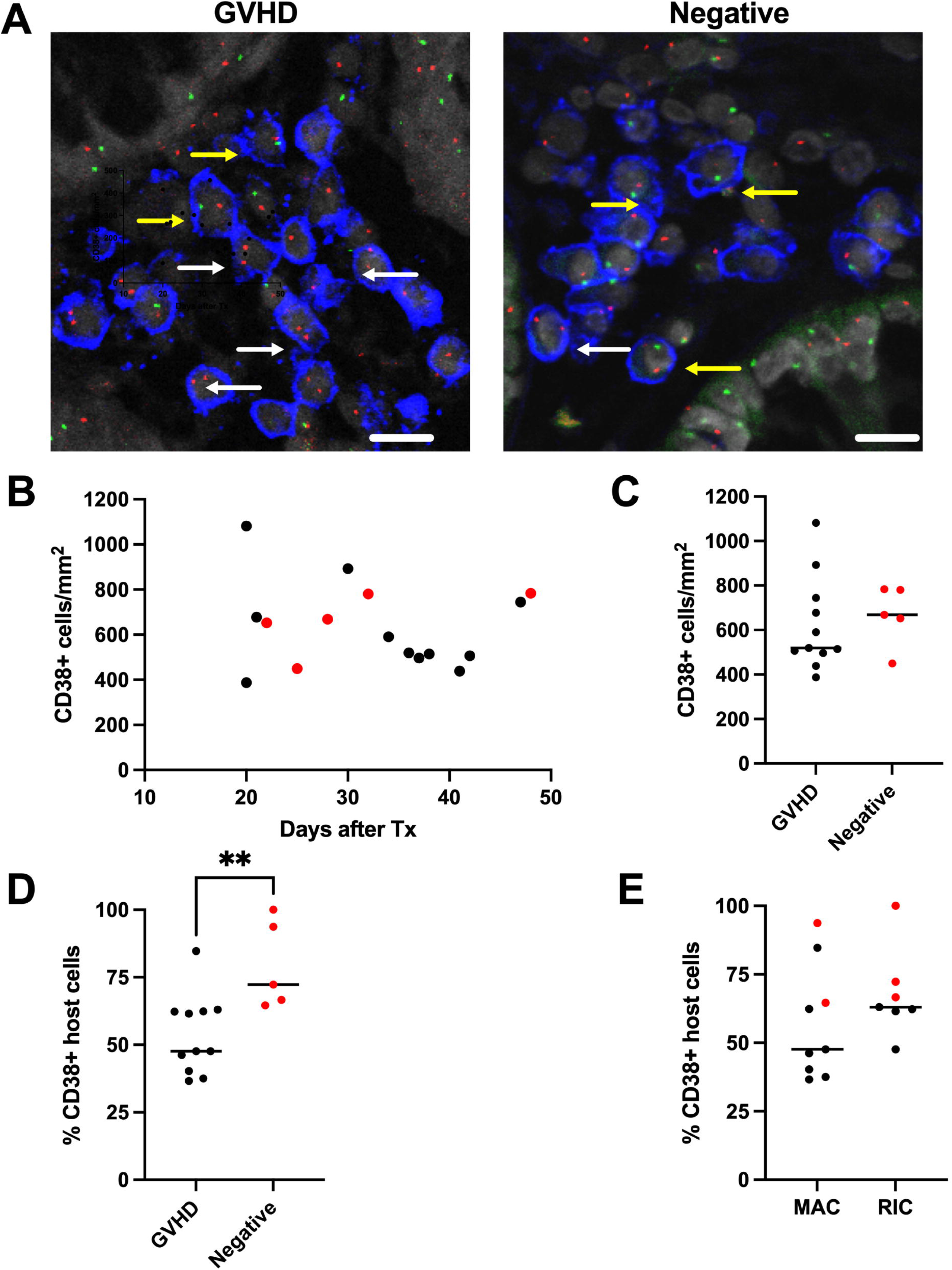
Colonic PCs are more rapidly replaced in acute GVHD. **(A)** Sections of FFPE colonic biopsies co-stained with X (red) and Y (green) FISH and anti-CD38 (blue) of male patients with (left) and without GVHD (right) receiving female donor graft. XX+ and XY+ PCs are arrowed (white and yellow, respectively). Scale bar 20 µm. **(B)** Density of CD38+ PCs in patients with (black) and without (red) GVHD related to time after transplantation (Tx) **(C)** Density of CD38+ PCs in patients with (black) and without (red) related to time after transplantation. **(D)** Percentage of host CD38+ PCs in patients with and without GVHD. ** p<0.0067. **(E)** Percentage of host CD38+ PCs in patients with (black) and without (red) GVHD receiving reduced-intensity conditioning (RIC) and myeloablative conditioning (MAC). Median values are indicated (horizontal lines).

**Table 1.** Clinical characteristics.

|  |  |
| --- | --- |
| **Number of patients** | 16 |
| **Age, median (range)** | 45 (21-72) |
| **Conditioning** |  |
| Myeloablative, n(%) | 11 (68) |
| Cy+Busulfan, n | 11 |
| Reduced intensity conditioning, n (%) | 5 (32) |
| Flu+Busulfan | 2 |
| Flu+Cy | 2 |
| Flu+Treo | 1 |
| **MHC allele match/mismatch** |  |
| Match, related | 8 (50) |
| Mismatch, related | 1 (6) |
| Matched, unrelated | 6 (38) |
| mismatched, unrelated | 1 (6) |
| **Stem cell source** |  |
| PBSCT, n (%) | 13 (81) |
| BMSCT, (%) | 3 (19) |
| **GVHD prophylaxis** |  |
| Siro+Ciclo, n (%) | 1 (6) |
| MTX+ Siro, n (%) | 15 (94) |
| **Underlying diagnosis** |  |
| AML, n (%) | 8 (50) |
| MDS, n (%) | 4 (25) |
| CMML, n (%) | 1 (6) |
| NHL, n (%) | 1 (6) |
| ALL, (%) | 2 (12) |

Cy: Cyclophosphamide, Flu: Fludarabine, Treo: Treosulfane
PBSCT: Peripheral blood stem cell, BMSCT: Bone marrow
Siro: Sirolimus, Ciclo: Ciclosporine, MTX: Tithotrexate
AML: Acute myeloid leukemia, MDS: Myelodysplasti syndrome
CMML: Chronic myelomonocytic leukemia, NHL: Non-Hodkins lymphoma
All: Acute lymphoblastic leukemia

## Discussion

Here we report that ultra long-lived PCs secret IgA antibodies that highly coated most bacterial taxa in ACHIM; a diverse anaerobic culture of human gut microbiome. IgA produced by long-lived as well as short-lived PCs also showed broad bacterial reactivity. There was a correlation between bacterial binding across the PC subsets.

We have previously reported that 60-70% of all SI PCs in adults have an average lifespan of more than 10 years and a significant fraction of such cells are likely to survive a lifetime^23^. Here we find that IgA produced by these ultra long-lived PCs are specific for the vast majority of taxa in a complex microbiota community. This shows that humoral memory responses to bacterial antigens in the human SI are extremely persistent. We believe that bacteria-specific secretory antibodies produced by ultra long-lived and long-lived PCs play an important role for long-term stability and resilience of the healthy intestinal microbiota community^24^.

It has been shown in a series of studies that patients with acute gut GVHD display low microbiota diversity^41,45,46^. The finding that intestinal host PCs were more rapidly replaced in allo-HCT patients with acute gut GVHD than those without acute gut GVHD further strengthened the notion that a stable humoral memory response is important for a diverse microbiota. The presence of long-lived PCs has been analyzed in two intestinal inflammatory disorders, celiac disease and ulcerative colitis; both associated with dysbiosis^47,48^. Reduced frequencies of ultra long-lived SI PCs were found in both treated and untreated celiac disease patients^49^ and the frequency of both ultra long-lived and long-lived PCs were reduced in active ulcerative colitis^50^. Together, these findings clearly link dysbiosis to reduced long lasting humoral memory.

*Hydrogenanaerobacterium* and *Prevotella* showed the highest enrichment score measuring the binding of IgA from ultra long-lived PCs. Importantly, *Prevotella* has been shown to have pro-inflammatory properties in IBD and decreased or increased levels of *Prevotella* are common features in several non-intestinal autoimmune diseases^51^. Our findings are in agreement with the notion that pathobionts are controlled by high binding of bacteria-reactive IgA. Moreover, *Blautia* and *Veilonella,* that have probiotic properties, showed low level IgA binding.

Ultra long-lived and long-lived SI PCs display high somatic hypermutation^23^, indicating that they develop through T-cell dependent pathways. We have recently shown that a high number of resident SI CD4 T cells also are long-lived^52^, suggesting that there is long-lasting T cell memory to gut antigens. Interestingly, in a mouse model it was recently shown that many TCR clones derived from the gut were specific to several bacterial strains. This “cross-strain” reactivity was shown to be caused by sharing of a conserved TCR epitope between strains^53^. This indicates that a limited number of T cell clones can provide T-cell help to a large number of B cell clones. Secretory IgA has broad reactivity with bacteria, most likely because of specific binding to bacterial glycan structures that are shared between taxa^54^. Together, this one-to-many TCR-to-strain relationship combined with cross-species reactivity of IgA suggest that interactions between T cells and B cells at mucosal surfaces provide a reasonable explanation for the high fraction of bacteria that is coated with secretory IgA. The gut microbiota has received enormous attention over the last years and their functions seems to be involved in a variety of disorders including neurodegenerative disorders, autoimmune diseases, cardiovascular disease, obesity and behavioral disorders^55^. Our findings suggest that ultra long-lived and long-lived PCs are directly involved the composition and functions of the gut microbiota and their potential role in microbiota-mediated health and disease should be studied further.

## Materials and Methods

### Human biological material

Duodenum-proximal jejunum tissue was resected from nonpathological SI during Whipple procedure (pancreaticoduodenectomy) of pancreatic cancer (n = 4; age range 54–82 yr) as described^23^. Briefly, the resected SI was opened longitudinally and rinsed thoroughly in PBS. Mucosal folds were dissected off the submucosa. To obtain single-cell suspensions for flow cytometry, cell sorting, and culture, epithelial cells were removed by shaking in PBS containing 2 mM EDTA three times for 20 min at 37°C, and the remaining lamina propria was minced and digested in RPMI medium containing 2.5 mg/ml Liberase and 20 U/ml DNase I (both from Roche) at 37°C for 1 h. Digested tissue was passed through 100-μm cell strainers (Falcon) and washed three times in PBS.

The study was approved by the Regional Committee for Medical Research Ethics in Southeast Norway and the Privacy Ombudsman for Research at Oslo University Hospital–Rikshospitalet and complies with the Declaration of Helsinki. All participants gave their written informed consent.

### Flow cytometry

Single-cell suspensions from lamina propria were stained on ice with anti-CD45 BV510, anti-CD19 PE/Cy7, anti-CD27 BV421, and anti-CD38 APC/Cy7 (all from Biolegend), and acquired on a BD LSRFortessa (BD Biosciences)Data were analyzed using FlowJo 10.

### Bead-based PC sorting for IgA harvesting

Single-cell suspensions from SI resections were stained with biotinylated (DSB-X Biotin Protein kit; ThermoFisher) anti-CD38 antibody (clone HB7 from Absolut Antibody LTd). Bead-free CD38+ cells were obtained after isolation with Flexicomp beads and subsequent incubation with releasing buffer, following the kit’s instructions (11061D; ThermoFisher). CD19+, CD45+, and CD45− PC subsets were magnetic-activated cell sorted in two sequential steps: first applying CD19 microbeads and, then, sorting the resulting negative fraction with CD45 microbeads (both from Miltenyi Biotec). Purity of sorted CD38+ PCs was assessed by flow cytometry and was consistently >80%.

### Cell culture and ELISA

Following sorting, PCs were cultured for 48 h in RPMI medium with 10% heat-inactivated fetal calf serum (FCS), l-glutamine and penicillin-streptomycin. Cells were spun down at 500 rcf for 7 min, and the supernatant was transferred to protein concentrator tube (Amicon Ultra). The supernatant was then diluted and transferred to 96-well plates precoated with rabbit anti–human IgA (A0262; Dako) and blocked with 1% BSA. Purified IgA monomer at known concentration was used as standard. Bound IgA was detected with goat anti–human IgA peroxidase conjugate (A0295; Sigma-Aldrich).

### IgA^+/-^ bacterial sort purification by FACS

Approximately 20x10^6^ bacteria from ACHIM (Anaerobically Cultivated Human Intestinal Microbiota) in a volume of 250 μl of PBS with 2% FCS containing 3.125 μg of antibodies harvested from the PC subsets. After 1 h of incubation at 4°C in a rotary shaker, 750 μl of flow buffer was added to the mixture and centrifuged at 10,000 g for 5 min. The centrifuged pellet was then stained with secondary antibody (anti-human IgA APC, Miltenyi Biotec) and incubated for 15 min at 4° C in a rotary shaker. After incubation the bacterial suspension was then washed and stained with 1μM SYTO BC green fluorescent nucleic acid stain (ThermoFisher). A minimum of 50,000 IgA^+^ or IgA-particles were acquired for each sample using a BD FACS Aria IIIu (BD Biosciences). Sorted samples were centrifuged at 10,000 g for 8 min at 4°C, resuspended in 20 μl Milli-Q water and stored at −80°C.

### 16S rRNA gene amplicon sequencing and analysis

The sorted bacterial samples were directly used as DNA template for the amplification of V3/V4 regions of 16S rRNA genes using primers 319F and 806R. PCR amplicons were normalized, pooled together and quality checked using Agilent Bioanalyzer 2100. FACS buffer collected during cell sorting were included in the library preparation as controls to identify potential contaminating sequencing reads. The pooled library were stored at −20°C until sequencing. The amplicon libraries were sequenced with single barcodes in paired-end mode (2 × 300 nt) using a MiSeq sequencer (Illumina). Paired-end reads were filtered for Illumina Universal Adapters, demultiplexed, quality trimmed. Denoising to ASVs, filtering of contaminants and rare ASVs were performed with QIIME2 and taxonomic classification are done with SILVA.

### Calculation for bacterial enrichment in IgA+ fraction

The enrichment score of IgA binding for each taxon was calculated as follows: Fraction of IgA+ bacteria/fraction of IgA-bacteria. An enrichment score of 1 means that there was a relative enrichment of the taxon in the IgA+ fraction. **Fluorescence in-situ hybridization (FISH) and immunofluorescence staining** Formalin-fixed and paraffin embedded (FFPE) tissue sections were cut at 5-6 µm and placed on superfrost glass slides and dried at 65 °C for 8-12 hrs. Slides were then deparaffinized in xylene 5% and rehydrated in ethanol. Then, slides were boiled in citrate buffer (pH=6.0) at 100°C for antigen retrieval and DNA denaturation. For sex-mismatched allo-HCT patients, FISH probes for X and Y chromosome (Abbot Molecular) were applied, and sections were incubated overnight at 37°C and then washed briefly in PBS and dH2O before staining with anti-human CD38 followed by secondary antibody The sections were counterstaining with Hoechst and mounted with Prolong Diamond (both from ThermoFisher).

### Image acquisition and analysis

Analysis of sections was performed as previously described (23). Briefly, slides were imaged at X200 magnification by confocal microscopy (Olympus FV1000) and further examined using ImageJ. Cell density is given as number of cells per mm2 of lamina propria. Area of lamina propria was determined by using the area measure tool in ImageJ. Host cells in male recipients with female donors were identified by the detection of chromosome Y. Because we examine section cut at 5-6 µm we are not able to identify both X and Y chromosomes in all cells. We have previously performed immune-staining experiments on sections from colon tissue of adult males with normal histology. By confocal microscopy (Olympus FV1000) we were able to detect positive Y signal in a median of 62% of immunostained cells^56^. Based on this result, we determined the density of host (male) cells in gender mismatched patients by multiplying Y+ immunostained cells with 1.6 (100/62= 1.6), and used this estimated density to calculate the percentage of host cells.

## Author contributions

E.S.B., N.B.P. and F.L.J. conceived the study. N.B.P., B.K.C., F.S., H.M.R., A.T.A. and T.G.D. contributed to study design and performed experiments. K.H. and D.D. contributed to data analysis. J.E.R.V., E.S.B. and F.L.J participated in data analysis and interpretation. N.B.P. and F.L.J. wrote the manuscript. All authors contributed to the editing of the manuscript.

