## Supplementary figures and images for "Ultra long-lived plasma cells in the human small intestine produce microbiota-reactive IgA antibodies"

### Supplemental file

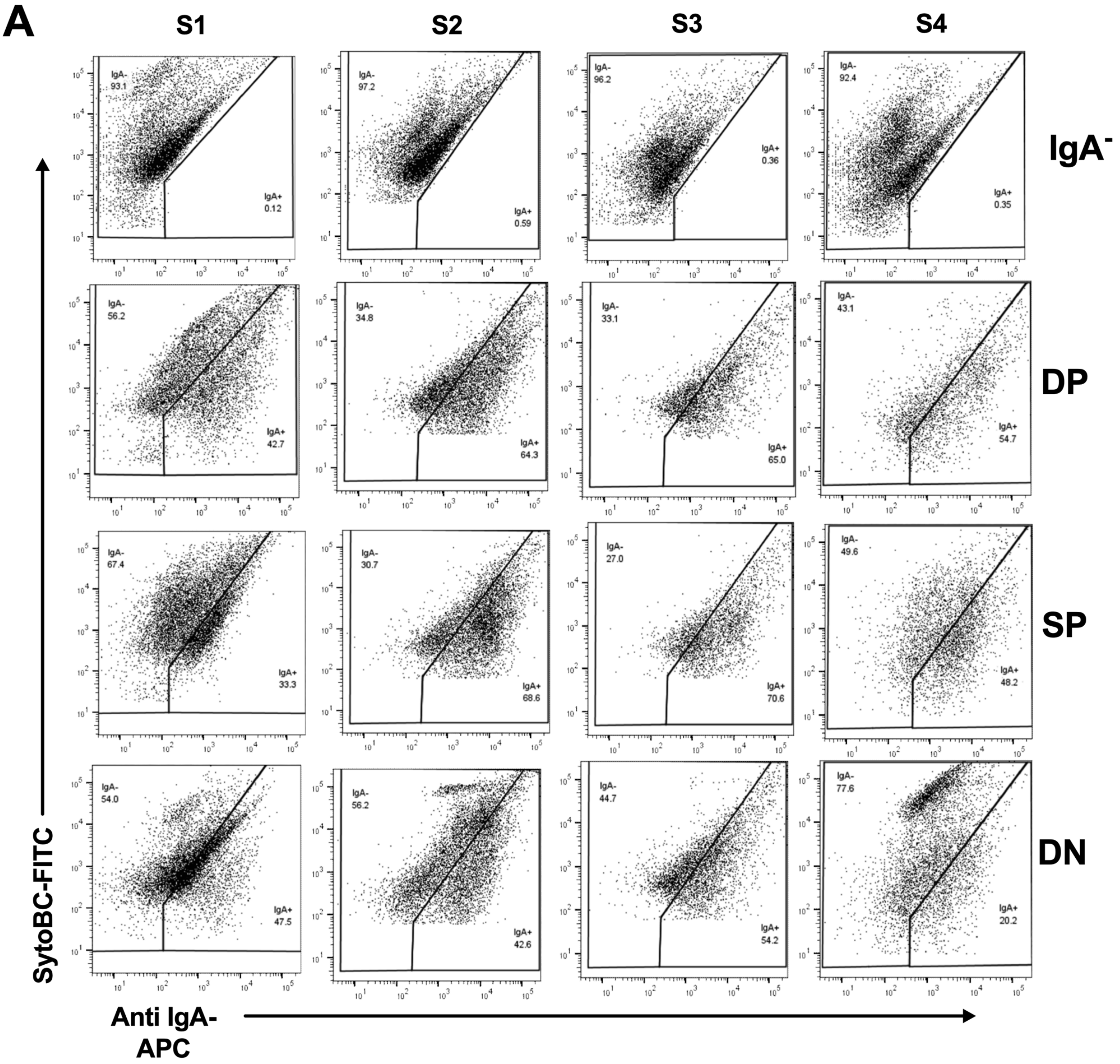

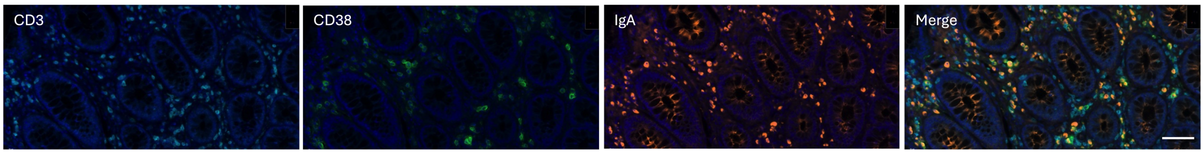
